# When Melodies Cue Memories: Electrophysiological Correlates of Autobiographically Salient Music Listening in Older Adults

**DOI:** 10.1101/2025.04.16.649223

**Authors:** Veronica Vuong, Mary J. O’Neil, Andrew Dimitrijevic, Bradley R. Buchsbaum, Michael H. Thaut, Claude Alain

## Abstract

Autobiographical memory is essential for older adults, providing a foundation for self-identity. Healthy aging is accompanied by changes in memory retrieval. However, musical memory remains relatively preserved, suggesting it may serve as an effective cue for autobiographical memory recall. Autobiographically salient (ABS) music (i.e., deeply encoded songs associated to important people, places, and events) is posited to engage distinct memory processes than familiar (FAM) music (i.e., songs that are recognized but lack personal significance). We tested this in 36 older adults (70.6 ± 6.6 years, 20 females) who listened to music of varying degrees of personal significance, including ABS, FAM, and unfamiliar (UFAM) music. In Experiment 1, participants pressed a button as quickly as possible when they recognized the excerpt as ABS, FAM, or UFAM. In Experiment 2, we measured event-related potentials and time-frequency responses while participants listened to the same stimuli and rated familiarity and memory at the end of each excerpt. Participants had the fastest reaction times for ABS, followed by FAM, then UFAM music. We observed a sustained evoked response from 2238 to 5000 ms post-stimulus onset that was largest in amplitude for ABS music, compared to FAM and UFAM music, over right frontal-central regions. We also observed less beta power suppression for ABS than FAM music between 1300 and 5000 ms over bilateral frontal-central-parietal areas. Our behavioral and neurophysiological findings show that ABS music is associated with faster and stronger memory-related activity distinct from FAM music.

## Introduction

Healthy aging is accompanied by well-documented changes in episodic autobiographical memory retrieval. Compared to young adults, older adults tend to recall episodic memories with fewer event-related details [1,2], less specificity [2,3] and diminished vividness [4]. Autobiographical memory is organized in a hierarchical model consisting of three levels with increasing specificity (see Conway and Pleydell-Pearce [5]). Successful retrieval can occur as a cascading effect, whereby activating higher-level representations facilitates access to event-specific details, leading to mental time travel [6]. Crucially, this process is influenced by the saliency of the cue.

Music can serve as a highly effective stimulus to trigger autobiographical memories [7]. For example, compared to words [8] and faces [9], music has been shown to elicit more episodic details. While the effectiveness of music as a memory cue may be attributed to the frequency with which it is listened to in daily life [10], the emotional quality it possesses [11], and/or associations to significant events [12], not all music is encoded into memory to the same degree. For instance, unfamiliar (UFAM) music refers to novel songs that we have not encountered before and, thus, neither recognize nor ascribe any personal significance to. Familiar (FAM) music includes those that we recognize through exposure but lack any personal meaning. Autobiographically salient (ABS) music is deeply encoded and is distinguished by its personal significance, evoking specific memories (i.e., a person, location, or event) and associated emotions. Remarkably, the capacity for ABS music to cue autobiographical memories persist in individuals with mild cognitive impairment and Alzheimer’s disease [13–16], underscoring music’s role as a bridge to the past. Understanding how the brain differentially engages with music conditions that vary by memory is pivotal to the development and implementation of effective music-based interventions for older adults, particularly those with dementia.

Given the distinct characteristics of ABS, FAM, and UFAM music, each type is expected to elicit different neural activity patterns, influenced by their varying degrees of personal significance. Previous neuroimaging research using functional magnetic resonance imaging (fMRI) and positron emission tomography (PET) have primarily focused on FAM and UFAM music [17]. Our Activation Likelihood Estimation meta-analysis of 23 neuroimaging studies showed that listening to FAM compared to UFAM music, or an equivalent control condition, yielded consistent activity in the left supplementary motor areas, left inferior frontal gyrus, and the left claustrum/insula [17]. Fewer studies have examined ABS relative to UFAM music, however, greater activation in the medial prefrontal cortex, precuneus, anterior insula, basal ganglia, hippocampus, amygdala, and cerebellum have been revealed [18]. Research that directly compares ABS and FAM music is lacking. Based on the above findings, different neural substrates may support the two music conditions.

In addition, the time needed to identify music excerpts may also vary. Prior studies used a gating paradigm where successive musical notes [19] or the duration of the excerpt [20–22] are gradually increased to assess the minimum amount of acoustic information (e.g., timbre or pitch) required to evoke a feeling of familiarity. Findings from these studies suggests that 200 to 500 ms in stimulus duration [20–22] is needed to elicit a feeling of familiarity but is insufficient to yield above-chance level identification of popular songs [23]. While a sense of familiarity may quickly emerge from hearing a musical except, the actual recollection of the corresponding artist and title requires more time. It is also unclear how much time listeners need to recognize musical excerpts associated with their autobiographical memory. Listeners may recognize ABS music more quickly due to deeper encoding. Moreover, the duration for familiarity and recognition above are based on young adult studies and may be longer in older adults due to age-related changes in sensory and cognitive processing.

Scalp-recordings of event-related potentials (ERPs) provide precise temporal information and can differentiate neural activity underlying familiarity and recollection during episodic memory retrieval [24]. For instance, correctly identified old from new items elicit more positive-going waveforms. In studies examining familiarity, this old/new effect has been observed over the mid-frontal scalp area, referred to as the FN400 (for a review, see Friedman and Johnson [25]). Recollection is indexed by an old/new effect over the left parietal scalp region between 400 and 800 ms post-stimulus onset, termed the Late Positive Component (LPC) [26]. A third relevant ERP modulation is a right-lateralized late frontal effect (LFE), emerging 600 to 2000 ms after stimulus onset [27]. The LFE is thought to index post-retrieval processing of contextual information [27,28] or self-relevant processing [29,30].

The above ERP components related to recognition memory have been observed in studies assessing musical memory in young adults. For instance, Jagiello et al. [31] reported an early ERP modulation from 350 to 750 ms post-stimulus onset over right frontal-central regions, which exhibited greater negativity for ABS than UFAM music. They also observed a subsequent ERP modulation, from 540 to 750 ms after music onset over the left parietal area, characterized by an increase in amplitude for ABS relative to UFAM music. A more recent study observed greater positivity for FAM than UFAM music from 400 to 450 ms over right and left frontal-central scalp regions [32]. This was accompanied by greater alpha power suppression when listening to FAM than UFAM music, over left frontal and posterior areas, along with low-beta suppression over frontal scalp areas. The former is interpreted as increased attentional engagement, the latter as reflecting familiarity-related processes. Although familiarity ratings were obtained, the possibility that some of the excerpts may have elicited an autobiographical memory cannot be excluded. The above studies shed light on the neural correlates of musical memory but were limited to only two experimental conditions. Thus, it remains unclear whether ABS music generates different neural responses than FAM music. A comparison of the neural responses to ABS, FAM, and UFAM music could provide deeper insight into the neural underpinnings of music perception and memory.

The present study used behavioral and neuroelectric methods to examine how healthy older adults process ABS, FAM, and UFAM music. In Experiment 1, we measured the time needed by older adults to recognize ABS, FAM, and UFAM music by asking participants to press a button as soon as they identified the excerpt as belonging to the respective music condition. Experiment 2 investigated ERPs and time-frequency responses across music conditions in the same group of participants. Familiarity and memory ratings were provided at the end of each excerpt to minimize the contamination of motor processes during music listening. We anticipated that the ERPs of the music conditions would differ in their sustained activity, demonstrating distinct memory processes associated with each contrast. Specifically, we hypothesized that the comparison between FAM and UFAM music would elicit a FN400 effect, reflecting increased familiarity-based processing. ABS music would produce a larger LPC when compared to UFAM music, indicating enhanced recollection-based retrieval, and a more pronounced LFE relative to FAM music, indexing greater self-referential processing or post-retrieval elaboration.

## Methods

### Participants

Participants were recruited if they were generally healthy, 60 years of age or older, English speaking, non-musicians (defined as not having acquired a Bachelor’s degree in music or equivalent, such as Royal Conservatory of Music [RCM] training at the Associate of the Royal Conservatory of Music [ARCT] level), attained a minimum of high school education, reported no diagnoses and/or hospitalizations due to head injury, depression, anxiety, psychiatric disorder, seizure disorder, alcohol or substance abuse, learning disabilities, tinnitus, major medical events/surgeries (e.g., open heart surgery), and brain radiation. All participants were screened to exclude medications affecting cognitive function and brain activity. Participants reported normal or corrected-to-normal vision and hearing.

Previous ERP studies showed differences between FAM and UFAM music [32] and ABS and UFAM [31] but used relatively small sample sizes (*n* = 20 and *n* = 10, respectively) that were limited to young adults. We aimed to recruit participants above these, due to the higher variability of older adults’ behavioral and brain responses [33,34]. Data were excluded from two participants due to non-compliance with the task(s), and two additional participants were excluded because of noisy electrophysiological data (e.g., excessive ocular and muscle artifacts). As a result, the final sample included 36 older adults (range = 61-86 years of age, 20 females. See Table 1 for demographic characteristics). All participants completed Experiment 1 and Experiment 2 on the same day.

**Table 1.** Demographic Characteristics (*n* = 36)

| Variable | Mean (*SD*) |
| --- | --- |
| Age (Years) | 70.6 (6.6) |
| Sex at Birth (F:M) | 20:16 |
| Handedness (R:L) | 34:2 |
| Bilateral Pure Tone Average Threshold (db HL) | 19.4 (9.6) |
| Montreal Cognitive Assessment | 28 (1.6) |
| Amount of Time Playing Instrument(s) (Years) | 4.2 (10.9) |

*F* = Female; *M* = Male; *R* = Right; *L* = Left; *db HL* = Decibels Hearing Level; *SD* = Standard Deviation.

This study was approved by the Research Ethics Board (REB) at the Rotman Research Institute, Baycrest Academy for Research and Education. Participants were recruited from the Rotman Research Institute participant database. All participants provided informed consent and received monetary compensation ($15/hour) plus transportation costs.

### Pre-Experiment Interview

We conducted an interview prior to the experimental sessions to identify 15 ABS musical titles with artist name(s) per participant, emphasizing that ABS music must bear personal significance (i.e., prompts a memory of a person(s), location, or event). Given that popular music, such as those played on the radio, typically contain vocals, and taking into consideration neural differences that may occur from listening to vocal vs. instrumental music, it was specified that all ABS songs must contain English vocals.

Using each ABS song, a FAM and UFAM song was selected and matched, to the best of our abilities, on the following aspects:

1. Artist style/genre.
2. Release year: Released the same or around the same year (± 5 years).
3. Number of online streams/plays: While we acknowledge that purchasing music albums was more commonly practiced among older adults, consistent data on album sales are not readily accessible. Online streaming is the dominant mode of music consumption. According to the Recording Industry Association of America [35], a song is certified Gold once it reaches 500,000 units (1 certification unit = 1 paid download or 150 streams), equating to 75 million plays across platforms. Spotify and YouTube are two of the largest streaming mediums and additionally provide play counts. We used a combined value of 50 million plays as a metric for FAM songs. Although this number is reduced, it does not include paid downloads or streams from other platforms such as Apple Music, Amazon Music, Tidal, Deezer, Pandora, and Bandcamp. All UFAM songs had fewer than 3 million combined plays on Spotify and YouTube. We selected the highest number of streams in the event of multiple YouTube videos and different metrics for the same song.

We utilized Spotify’s recommendation algorithm, which suggests songs with similar attributes. We refrained from selecting FAM and UFAM matches by the same ABS artist(s). All selected songs were reviewed by a musician on the study team to ensure that the songs matched appropriately. Although we did not initially choose FAM songs based on whether they were on the Billboard charts, we recognize that this metric might be more relevant for older songs (e.g., from the 1940’s or 1950’s), which precede the development of streaming platforms. We subsequently collected this data for each song (see Table 1 in the Supplementary Material for examples).

### Experiment 1: Behavioral Study Materials & Methods

#### Stimuli

Stimuli were music excerpts from the participant’s playlist (i.e., ABS music) and matched FAM and UFAM music. Each excerpt was trimmed to 10 seconds at a salient part of the song (e.g., the chorus) using Audacity® (version 3.2.2) [36]. Then, using a custom script in Praat (version 6.4.05) [37], the mean intensity was scaled to 70 dB SPL. An acoustic analysis (not shown) revealed that the excerpts were broadly comparable across conditions on properties including root mean square intensity, tempo, and zero crossing rate. While significant differences were observed in onset rise time, pitch, spectral centroid and spectral bandwidth, these were relatively modest in magnitude.

#### Procedure

First, participants familiarized themselves with the task by completing four practice trials with excerpts not used in the experimental run. Then, participants completed five blocks of 45 trials each (15 trials per music type). Thus, each excerpt was presented five times. Within each block, the excerpts from the ABS, FAM, and UFAM conditions were presented randomly. Music excerpts were presented binaurally via insert earphones at 55 db SPL.

On each trial, participants heard an excerpt from one of three conditions and indicated as quickly and accurately as possible whether it belonged to the ABS, FAM, or UFAM category by pressing one of three keys on a computer keyboard. ABS music was specified as personally significant; FAM music was indicated as music that was recognized but lacked any personal significance; and UFAM music was defined as music that had not been heard before. We emphasized that these categorizations of music were to be framed within their lifetime (i.e., hearing a UFAM clip for the second time does not qualify as a familiar stimulus and, therefore, would still be considered as belonging to the UFAM condition). Once the key was pressed, the music stopped, and the subsequent trial began three seconds later. The button assignment was counterbalanced across participants, resulting in six possible configurations. No feedback was provided after each trial or block. The experiment was programmed using Presentation software (version 23.0) [38].

#### Statistical Analysis Behavioral Analysis

A repeated-measures ANOVA was performed separately for accuracy and RT from correct responses, with music condition as the within-subjects factor, using R software (version 2023.12.1+402) [39]. Sphericity was tested using Mauchly’s test (*W*). When applicable, the degrees of freedom were adjusted using Greenhouse-Geiser (*ε*) estimates. Bonferroni corrections were applied for pairwise comparisons and are reported herein.

## Results

### Behavioral Results

Figure 1a and 1b show the group average accuracy and RT for each music condition. The repeated-measures ANOVA on accuracy yielded a main effect of condition (*F*_(2,70)_ = 24.90, *p* < 0.001, *η*^2^_p_ = 0.27). Pairwise comparisons revealed significantly higher correct responses for ABS music (*M* = 95.5%) than FAM (*M* = 76.8%) and UFAM (*M* = 78.7%) music (*p* < 0.001 in both cases). There was no difference in mean correct responses between FAM and UFAM conditions (*p* = 1). The percentage of trials in which ABS music was misidentified as FAM and UFAM music was 3.6% and 0.8%, respectively. Participants incorrectly identified FAM music as ABS and UFAM music in 10% and 13.1% of trials. Lastly, in trials involving UFAM music, 20.1% and 1.3% of trials were incorrectly identified as FAM and ABS music, respectively.

**Fig. 1.**
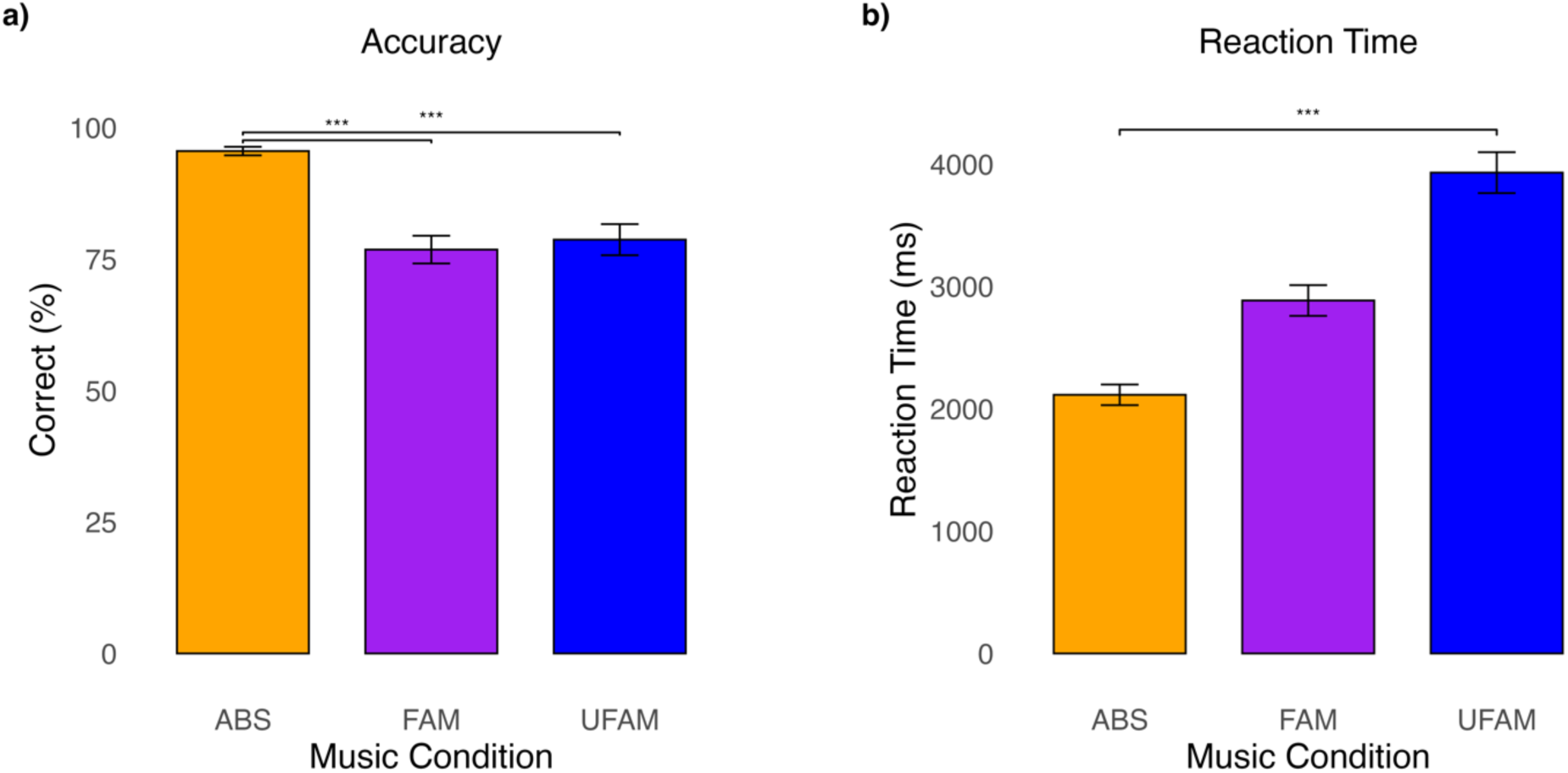
**a)** Mean accuracy and **b)** Mean reaction time (based on correct responses) by condition. *ABS* = Autobiographically salient; *FAM* = Familiar; *UFAM* = Unfamiliar Error bars indicate the standard error of the mean (SEM). *** *p* < 0.001

The repeated-measures ANOVA also revealed a main effect of condition on RT (*F*_(2,70)_ = 92.95, *p* < 0.001, η^2^_p_ = 0.48). ABS music was identified the fastest (*M* = 2114 ms, *SD* = 506 ms), intermediate for FAM music (*M* = 2885 ms, *SD* = 754 ms), and slowest for UFAM music (*M* = 3930 ms, *SD* = 1003 ms). Pairwise comparisons showed that all music conditions significantly differed from each other (*p* < 0.001 for all comparisons).

### Experiment 2: EEG Study Materials & Methods Stimuli

For each participant, the same set of music excerpts from Experiment 1 was used, with the only difference being that they were shortened to five seconds using Audacity (version 3.2.2) [36]. This duration was motivated by the findings from Experiment 1 and aimed to increase the number of trials for the ERP analyses.

#### Procedure

Participants were instructed to avoid movement as much as possible during EEG recording. First, two practice trials with music not used in the experimental run were presented. Then, participants completed five blocks of 45 trials each, in which the ABS, FAM, and UFAM excerpts were presented in a random order, with an inter-trial interval of three seconds. The music excerpts were presented binaurally through insert earphones at the same intensity as in Experiment 1. During music listening, participants were asked to direct their gaze on a white fixation cross in the center of a black screen. After each music excerpt, participants used a 4-point scale to rate familiarity and strength of memory association (1 = No familiarity/memory association; 2 = Some familiarity/memory association; 3 = Moderate familiarity/memory association; 4 = High familiarity/memory association) using the keyboard. The ratings were self-paced so that once the memory rating was complete, the next trial was initiated.

#### Electroencephalography Acquisition and Processing

Neuroelectric brain activity was recorded continuously using a 64-channel BioSemi ActiveTwo acquisition system and digitized at a sampling rate of 2048 Hz with a bandpass of DC-100Hz. The electrodes were positioned on the scalp according to the standard 10-20 system with a Common Mode Sense active and a Driven Right Leg passive electrode serving as the ground. Voltage offsets were kept below ± 40 mV.

EEG data were processed offline using Brain Electrical Source Analysis (BESA) Research software (version 7.1). Defective electrodes were interpolated using values from surrounding electrodes. Up to seven electrodes were interpolated per participant. Artifacts from ocular movements were identified from the recording and used to generate spatial components that model ocular artifacts, such as eye blinks as well as vertical and lateral eye movements. The spatial topographies were then subtracted from the continuous EEG to correct for eye movements and eye blinks. The data were downsampled to 256 Hz, re-referenced to the average of all electrodes, and digitally filtered with a 0.01 Hz high pass filter (forward phase, 6 dB/octave) and a 30 Hz low pass filter (zero phase, 24 dB/octave). Data were segmented into epochs, including 500 ms pre-stimulus and 6000 ms post-stimulus activity. The EEG recordings were scanned for artifacts. Trials contaminated by excessive peak-to-peak deflection (120 µV) were marked and excluded from the analysis. On average, 95.13% of trials were accepted for each music condition. The analysis epochs were averaged and baseline-corrected with respect to a 500 ms pre-stimulus interval.

#### Time-Frequency Analysis

The continuous EEG data were decomposed into time-frequency representation using complex demodulation with a 100 ms time window and 0.5 Hz frequency resolution from 1 to 30 Hz. Baseline correction was applied using a pre-stimulus interval from -500 to 0 ms.

#### Statistical Analyses Behavioral Data Analysis

A one-way repeated-measures ANOVA was performed separately to assess the effects of music conditions on familiarity and memory ratings. A Greenhouse-Geisser correction was used when the assumption of sphericity was violated. Planned pairwise comparisons were conducted using Bonferroni correction to examine significant main effects. Statistical analyses were performed using R software (version 2023.12.1+402) [39].

#### ERP and Time-Frequency Analyses

ERP waveforms and time-frequency representations were analyzed using a within-group ANOVA in BESA Statistics software (version 2.1). This software identifies clusters in time and space and uses a series of F-tests to compare the ERP amplitude between experimental music conditions at every time point. Preliminary clusters were identified in time (adjacent time points) and space (adjacent electrodes) where brain responses differed between the experimental conditions. The neighbor distance (circle arc length between two points) was set to 4 cm and the significance level of cluster building was set to 0.05, with 5000 permutations. We examined the brain responses (ERPs, time-frequency representations) across conditions from 0 to 5000 ms. Only significant clusters with at least four channels were included.

#### Brain-Behavior Correlation Analyses

Correlation analyses were computed between neuroelectric measures including ERP amplitude, alpha power, and beta power and behavioral measures including RT, familiarity, and memory ratings, across participants. Using BESA Statistics (version 2.1), a time window of 0 to 5000 ms was specified. The correlation clusters were corrected for multiple comparisons using Monte-Carlo resampling techniques [40], with a *p*-value of 0.05 and 5000 permutations. Similar to the ERP and time-frequency analyses, only significant clusters with at least four channels were included.

## Results

### Behavioral Data Results

Figures 2a and 2b show the group mean familiarity and memory rating as a function of music condition. The one-way repeated-measures ANOVA yielded a main effect of condition on mean familiarity ratings (*F*_(2,70)_ = 467.18, *p* < 0.001, η^2^_p_ = 0.881). Familiarity ratings were highest for ABS music (*M* = 3.95, *SD* = 0.09), followed by FAM music (*M* = 3.17, *SD* = 0.51), and lowest for UFAM music (*M* = 1.47, *SD* = 0.41) (all pairwise comparisons *p* < 0.001).

**Fig. 2.**
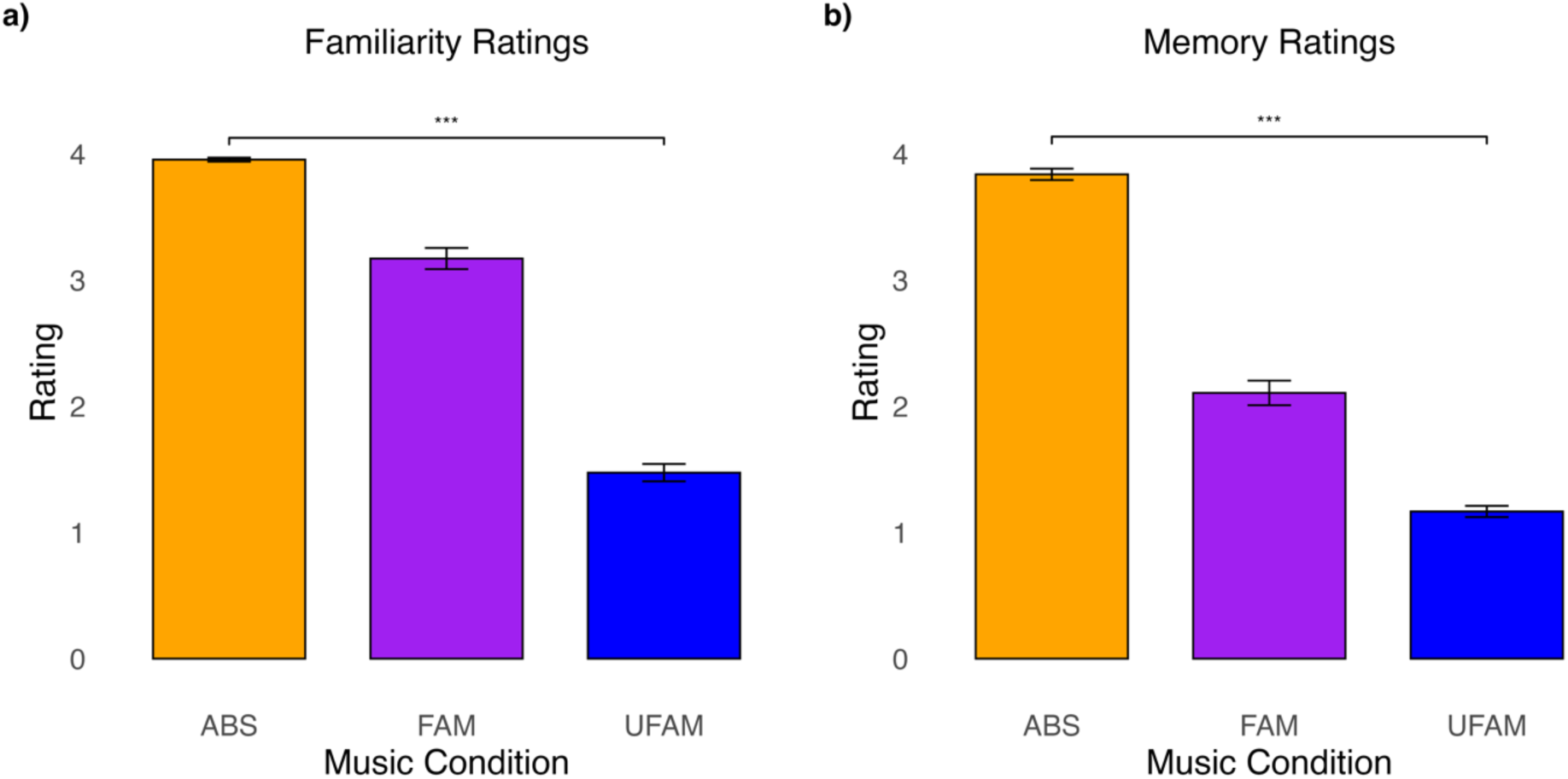
**a)** Mean familiarity and **b)** Mean memory ratings by music condition *ABS* = Autobiographically salient; *FAM* = Familiar; *UFAM* = Unfamiliar. Error bars indicate the standard error of the mean (SEM). *** *p* < 0.001

A one-way repeated-measures ANOVA on memory rating also showed a main effect of condition (*F*_(2,70)_ = 533.36, *p* < 0.001, η^2^_p_ = 0.883), with the highest memory ratings for ABS music (*M* = 3.83, *SD* = 0.27), then FAM music (*M* = 2.10, *SD* = 0.59), and lastly, UFAM music (*M* = 1.16, *SD* = 0.26) (Figure 2b) (all pairwise comparisons *p* < 0.001).

### ERP Results

Figure 3a and 3b shows the group mean ERP waveforms and topographies elicited by ABS, FAM, and UFAM music, which were time-locked to the onset of the music excerpt. All music conditions generated transient onset responses that inverted polarity between frontal-central and temporal-parietal electrodes, consistent with neural generators in the planum temporale and Heschl’s gyrus along the superior temporal gyrus [41,42]. These transient responses were followed by a large positive wave over the parietal region, a sustained evoked potential largest over right frontal-central, and offset responses that were largest over midline central areas.

**Fig. 3.**
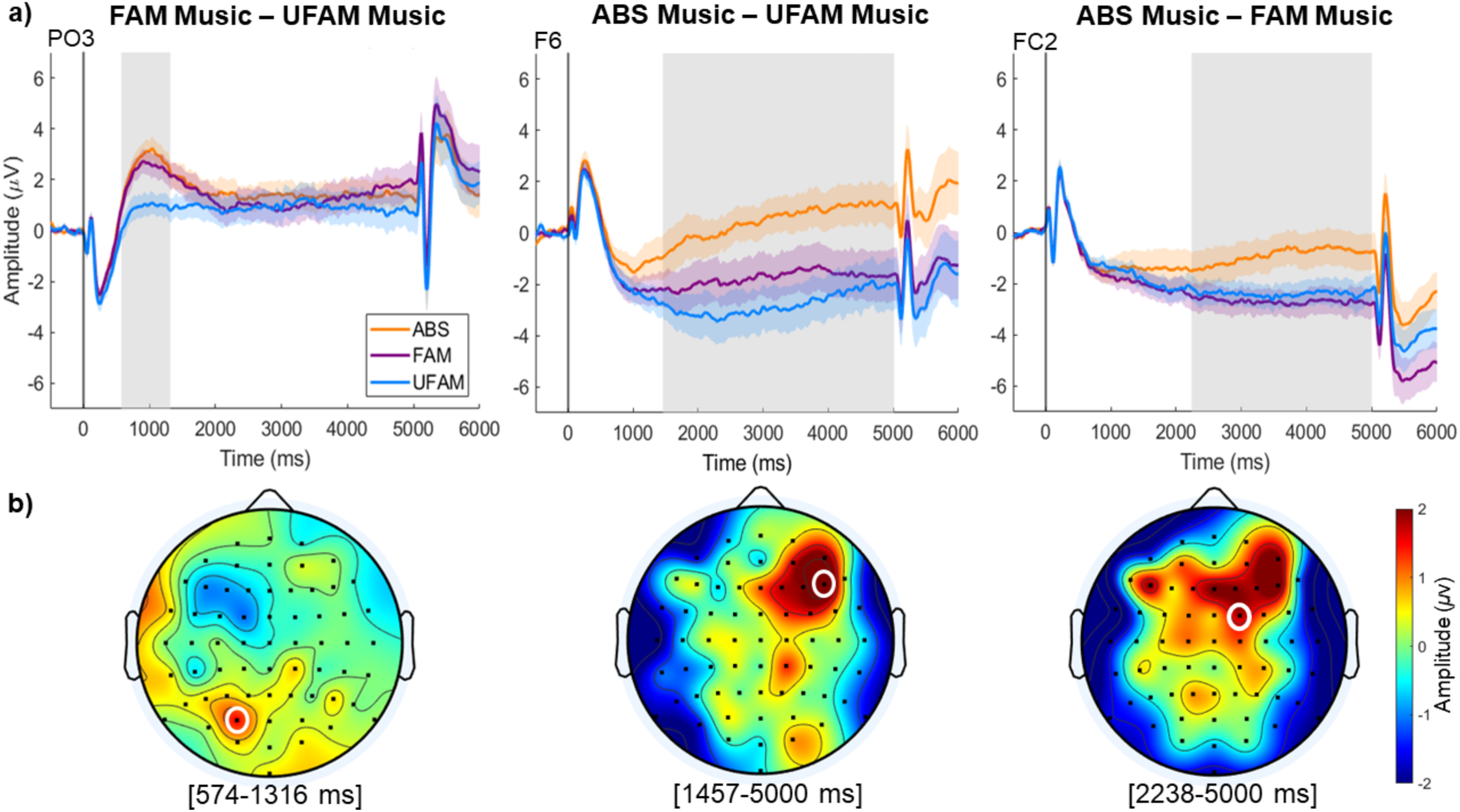
**a)** Grand averaged waveforms recorded at left parietal-occipital (PO3), right frontal (F6) and right frontal-central (FC1) electrodes, elicited by ABS music (orange), FAM music (purple), and UFAM music (blue), time-locked to the onset of music stimuli. Shading around each line represents the mean ± SEM. The gray box represents the time window where event-related potential (ERP) amplitudes significantly differed between planned pairwise contrasts. Positive voltage is plotted upwards **b)** Topographic maps illustrating the mean ERP amplitude difference between each planned pairwise contrast across the corresponding time windows of significance. The white circle indicates the electrode at maximum amplitude. The color bar represents the amplitude. *ABS* = Autobiographically salient; *FAM* = Familiar; *UFAM* = Unfamiliar.

When examining evoked activity from 0 to 5000 ms, cluster-based statistics revealed two significant clusters for the FAM and UFAM pairwise contrast. The first cluster was observed between 574 and 1316 ms, whereby FAM music was more positive in amplitude than UFAM music over left parietal-occipital electrodes, similar to an LPC. The second cluster, representing a polarity reversal, was observed between 770 and 1637 ms over left frontal-central regions.

The ABS and UFAM pairwise contrast indicated four significant clusters, with the most significant cluster observed between 1457 and 5000 ms (*p* < 0.001) over the right frontal cortex. The waveform for ABS music was more positive-going than for UFAM music.

Lastly, the ABS and FAM pairwise contrast yielded two significant clusters. The first cluster exhibited the most prominent effect, from 2238 to 5000 ms post-stimulus onset (*p* < 0.001), indicating more positive sustained ERP amplitude when listening to ABS than FAM music. This modulation was largest over the right frontal-central region. For a detailed list of all clusters, see Table 2.

**Table 2.**
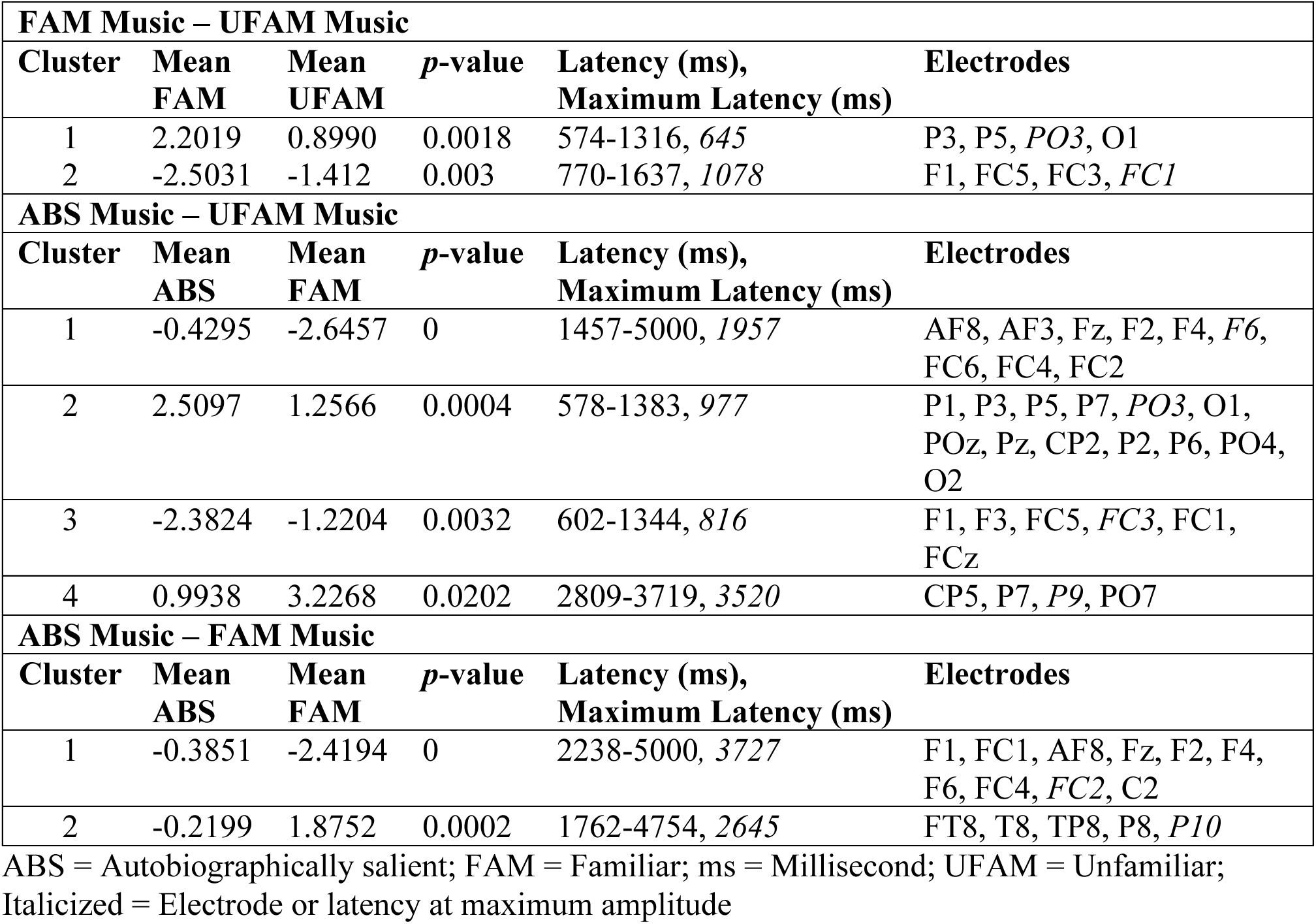
Summary of Cluster-Based Permutation Statistics from ERP Analysis.

| **FAM Music – UFAM Music** |  |  |  |  |  |
| --- | --- | --- | --- | --- | --- |
| **Cluster** | **Mean FAM** | **Mean UFAM** | ***p*-value** | **Latency (ms),**<br>**Maximum Latency (ms)** | **Electrodes** |
| 1 | 2.2019 | 0.8990 | 0.0018 | 574-1316, 645 | P3, P5, *PO3*, O1 |
| 2 | -2.5031 | -1.412 | 0.003 | 770-1637, 1078 | F1, FC5, FC3, *FC1* |
| **ABS Music – UFAM Music** |  |  |  |  |  |
| **Cluster** | **Mean ABS** | **Mean FAM** | ***p*-value** | **Latency (ms),**<br>**Maximum Latency (ms)** | **Electrodes** |
| 1 | -0.4295 | -2.6457 | 0 | 1457-5000, 1957 | AF8, AF3, Fz, F2, F4, *F6*, FC6, FC4, FC2 |
| 2 | 2.5097 | 1.2566 | 0.0004 | 578-1383, 977 | P1, P3, P5, P7, *PO3*, O1, POz, Pz, CP2, P2, P6, PO4, O2 |
| 3 | -2.3824 | -1.2204 | 0.0032 | 602-1344, 816 | F1, F3, FC5, *FC3*, FC1, FCz |
| 4 | 0.9938 | 3.2268 | 0.0202 | 2809-3719, 3520 | CP5, P7, *P9*, PO7 |
| **ABS Music – FAM Music** |  |  |  |  |  |
| **Cluster** | **Mean ABS** | **Mean FAM** | ***p*-value** | **Latency (ms),**<br>**Maximum Latency (ms)** | **Electrodes** |
| 1 | -0.3851 | -2.4194 | 0 | 2238-5000, 3727 | F1, FC1, AF8, Fz, F2, F4, F6, FC4, *FC2*, C2 |
| 2 | -0.2199 | 1.8752 | 0.0002 | 1762-4754, 2645 | FT8, T8, TP8, P8, *P10* |

ABS = Autobiographically salient; FAM = Familiar; ms = Millisecond; UFAM = Unfamiliar; Italicized = Electrode or latency at maximum amplitude

### Time-Frequency Results

Figure 4 shows the group mean temporal spectral evolution (TSE) spectrograms for the three conditions and specific contrasts between FAM and UFAM and ABS and FAM conditions. For all condition-specific spectrograms, transient theta power can be observed at music onset and offset. In addition, FAM and UFAM Music showed alpha and beta synchronization between 500 and 1500 ms after stimulus onset.

**Fig. 4.**
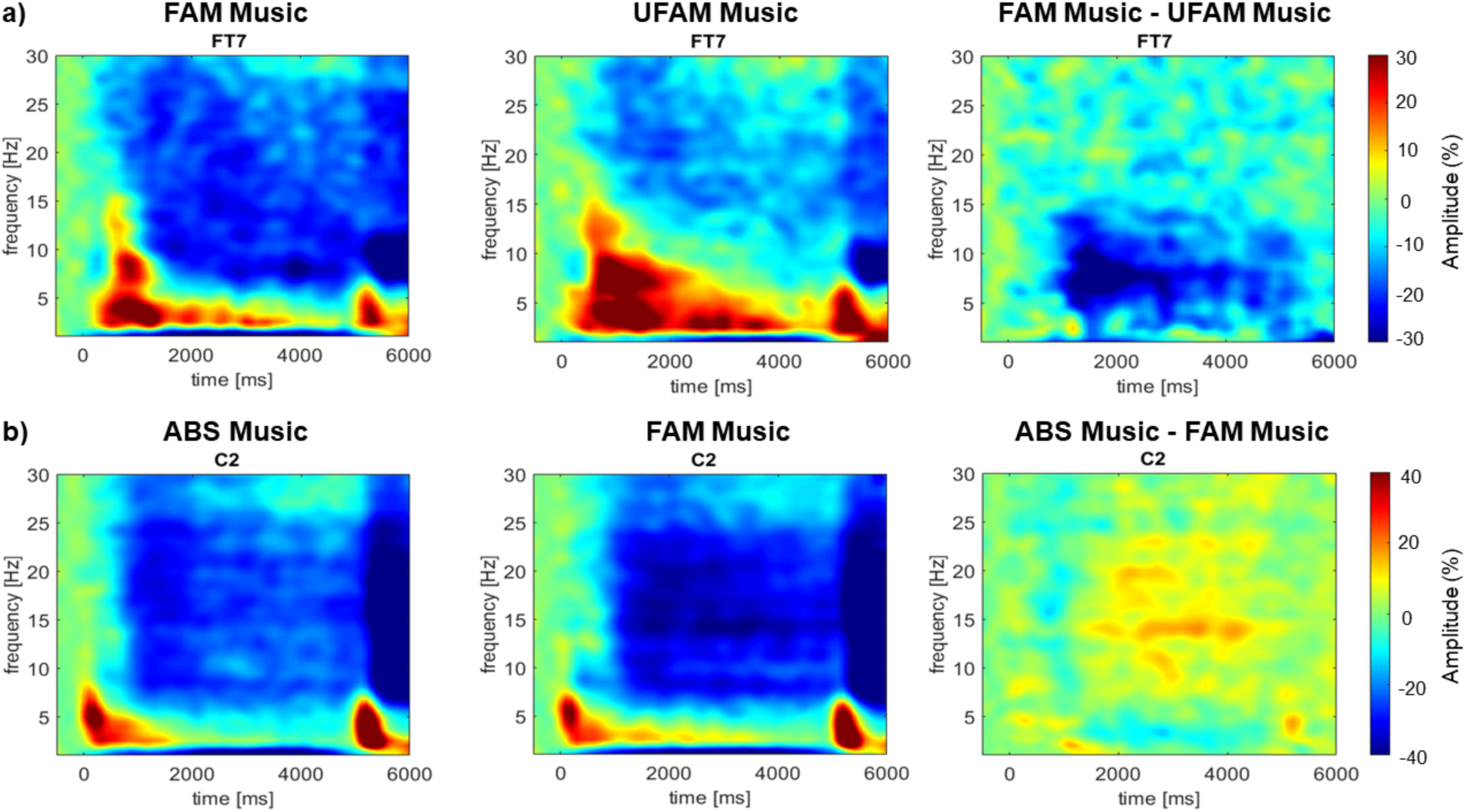
**a)** Group mean time-frequency representations over the left frontal-temporal (FT7) and **b)** midline-right central region (C2). Time 0 indicates stimulus onset. The color scale represents percent power change relative to the baseline. Greater alpha and low-beta suppression is observed when listening to FAM compared to UFAM music and ABS music. *ABS* = Autobiographically salient; *FAM* = Familiar; *UFAM* = Unfamiliar

When comparing FAM and UFAM music, the cluster analysis procedure and permutation-based statistics revealed one significant spatiotemporal cluster (*p* < 0.001). The cluster started at stimulus onset and ended at 5000 ms, encompassing delta, theta, alpha, and beta bands. It peaked at 1800 ms and was maximum in the alpha band (11 Hz) over the left frontal-temporal scalp area (FT7), revealing greater suppression for FAM music than UFAM music (Fig. 4A).

For the ABS and UFAM music contrast (not shown), two significant clusters were identified. The most significant cluster (*p* < 0.001) spanned the entire epoch and encompassed all frequencies, with a peak in alpha power (11.5 Hz) over the left central scalp area (C1). Greater suppression for ABS music relative to UFAM music was observed. As this contrast was outside our interest, the time-frequency response is not shown here.

The ABS and FAM music contrast revealed three significant clusters. The primary significant cluster (*p* < 0.001) began at 1200 ms after stimulus onset and lasted until the end of the music excerpt, extending across theta, alpha, and beta bands. The maximum difference occurred in the beta band (14 Hz) at 3400 ms over the right central scalp area (C2), indicating greater suppression for FAM music than ABS music (Fig. 4B).

Previous research showed greater suppression of alpha and low-beta power when listening to FAM than UFAM music [32]. Based on these findings, we compared alpha (8 Hz to 12 Hz) and low-beta (12 Hz to 16 Hz) bands across conditions.

### Frequency Band Specific Analysis

#### Alpha Frequency Band

Figure 5a shows the temporal dynamics of alpha power activity, averaged across electrodes identified from cluster-based permutation statistics.

**Fig. 5.**
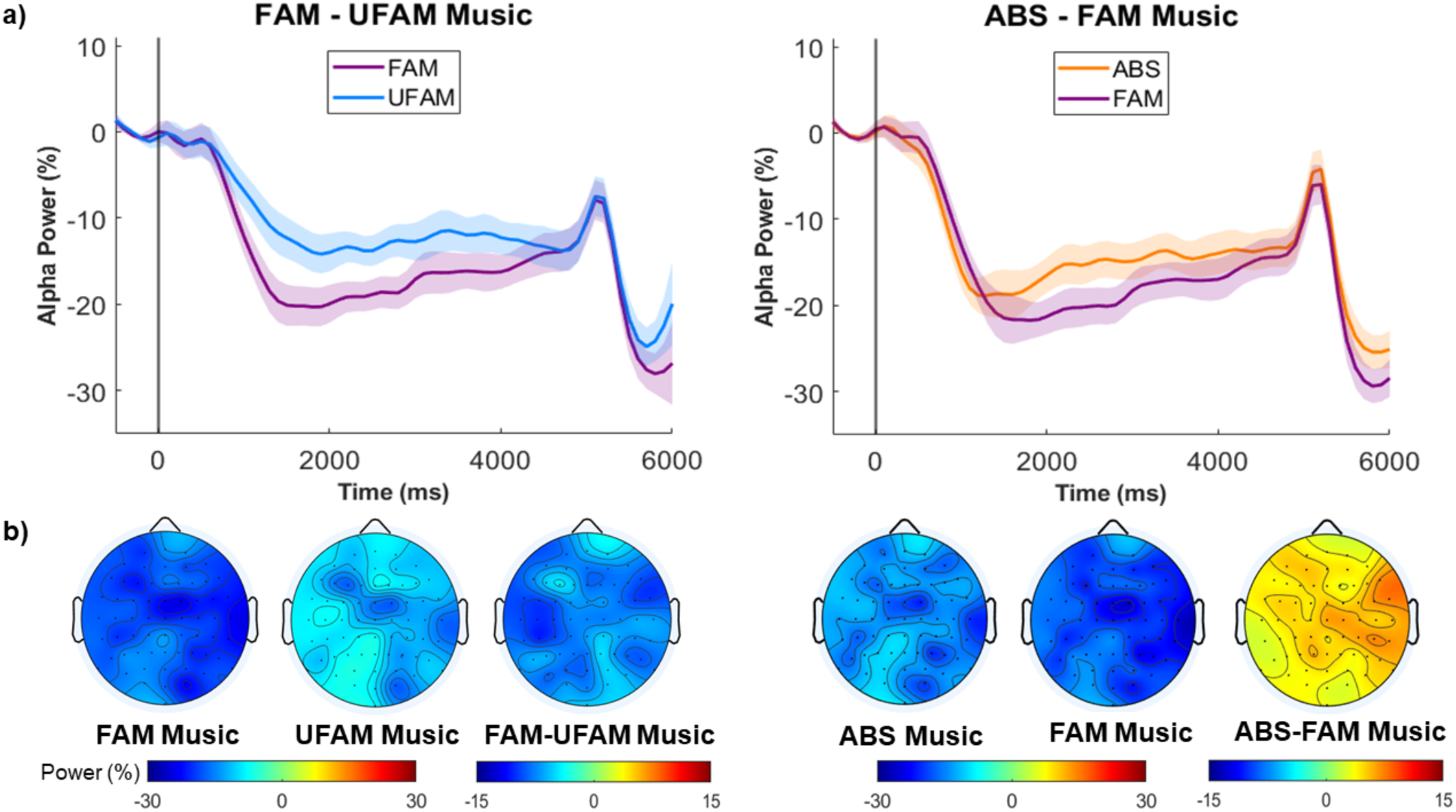
**a)** Changes in alpha power (8 to 12 Hz) relative to the pre-stimulus baseline, averaged over electrodes that are encompassed in each respective cluster. Greater alpha suppression is observed for FAM music compared to UFAM music (left panel) and ABS music (right panel). The amplitudes of each music condition are represented by different colors, where UFAM is blue, FAM is purple, and ABS is orange. Time 0 indicates the onset of the stimuli. Shading around each line represents the mean ± SEM. *ABS* = Autobiographically salient; *FAM* = Familiar; *UFAM* = Unfamiliar **b)** Topographic maps of the average of all channels, around the peak latency (+/-100 ms) for each pairwise contrast including FAM-UFAM (left panel) and ABS-FAM music (right panel)

When examining changes in alpha power over time, one cluster was observed for the FAM-UFAM music contrast, revealing a reduction in alpha power for the former condition compared to the latter (*p* < 0.001). This cluster was widely distributed over bilateral frontal-central-parietal regions (Table 3) and occurred between 300 and 5000 ms with the peak occurring at 1300 ms (Fig. 5a, left panel).

**Table 3.**
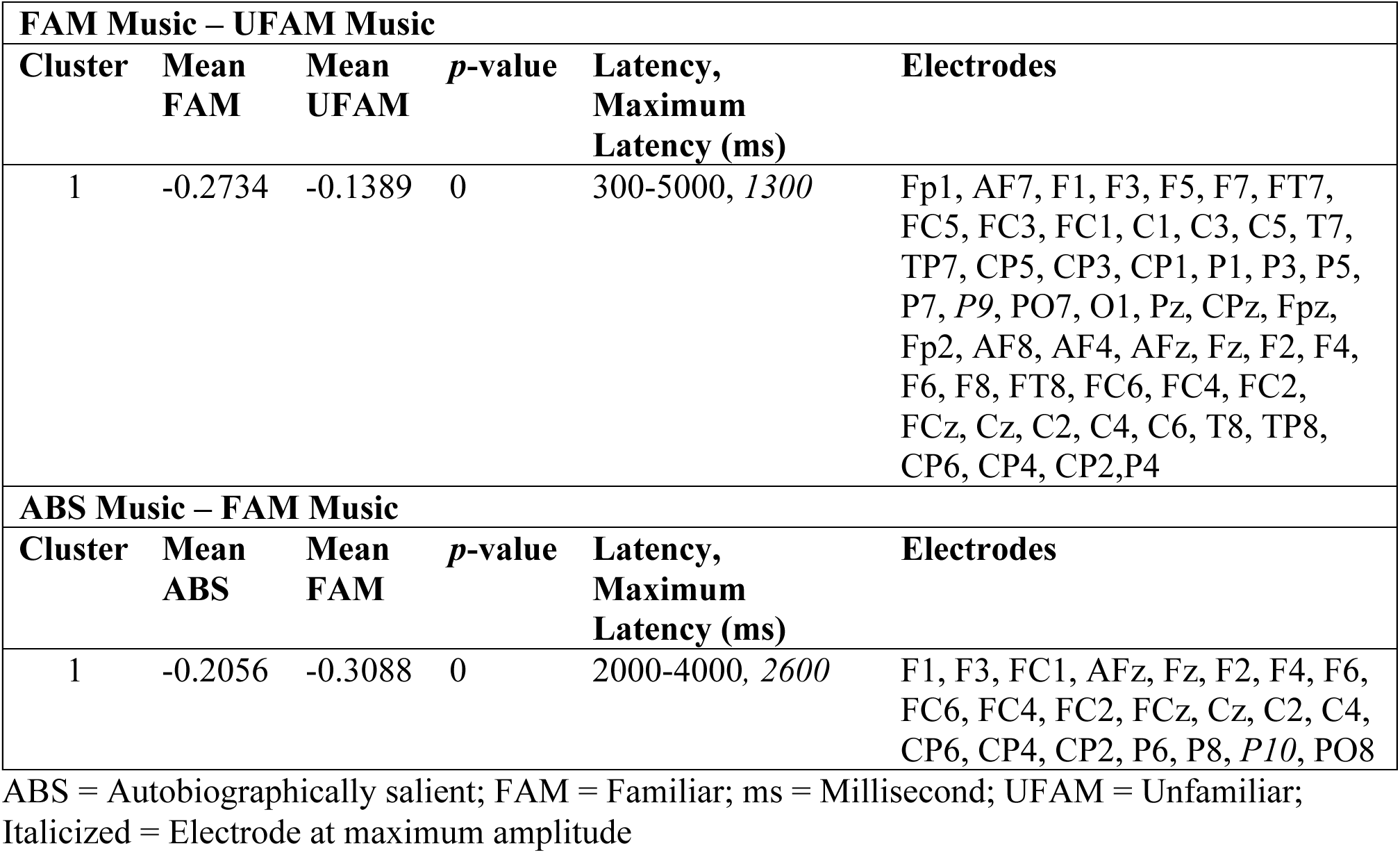
Summary of Cluster-Based Permutation Statistics from Alpha Band Power Analysis.

| **FAM Music – UFAM Music** |  |  |  |  |  |
| --- | --- | --- | --- | --- | --- |
| **Cluster** | **Mean FAM** | **Mean UFAM** | ***p*-value** | **Latency, Maximum Latency (ms)** | **Electrodes** |
| 1 | -0.2734 | -0.1389 | 0 | 300-5000, *1300* | Fp1, AF7, F1, F3, F5, F7, FT7, FC5, FC3, FC1, C1, C3, C5, T7, TP7, CP5, CP3, CP1, P1, P3, P5, P7, *P9*, PO7, O1, Pz, CPz, Fpz, Fp2, AF8, AF4, AFz, Fz, F2, F4, F6, F8, FT8, FC6, FC4, FC2, FCz, Cz, C2, C4, C6, T8, TP8, CP6, CP4, CP2, *P4* |
| **ABS Music – FAM Music** |  |  |  |  |  |
| **Cluster** | **Mean ABS** | **Mean FAM** | ***p*-value** | **Latency, Maximum Latency (ms)** | **Electrodes** |
| 1 | -0.2056 | -0.3088 | 0 | 2000-4000, *2600* | F1, F3, FC1, AFz, Fz, F2, F4, F6, FC6, FC4, FC2, FCz, Cz, C2, C4, CP6, CP4, CP2, P6, P8, *P10*, PO8 |

ABS = Autobiographically salient; FAM = Familiar; ms = Millisecond; UFAM = Unfamiliar; Italicized = Electrode at maximum amplitude

The ABS-FAM contrast also revealed one significant cluster indicating greater alpha suppression for FAM music relative to ABS music (*p* < 0.001). The effect occurred later, between 2000 and 4000 ms, peaking at 2600 ms (Fig. 5a, right panel) and was right-lateralized over frontal-central-parietal areas. For details on the cluster-based permutation statistics, see Table 3.

#### Low-Beta Frequency Band

Figure 6a shows the temporal dynamics of alpha power activity, averaged across electrodes identified from cluster-based permutation statistics for FAM-UFAM and ABS-FAM music.

**Fig. 6.**
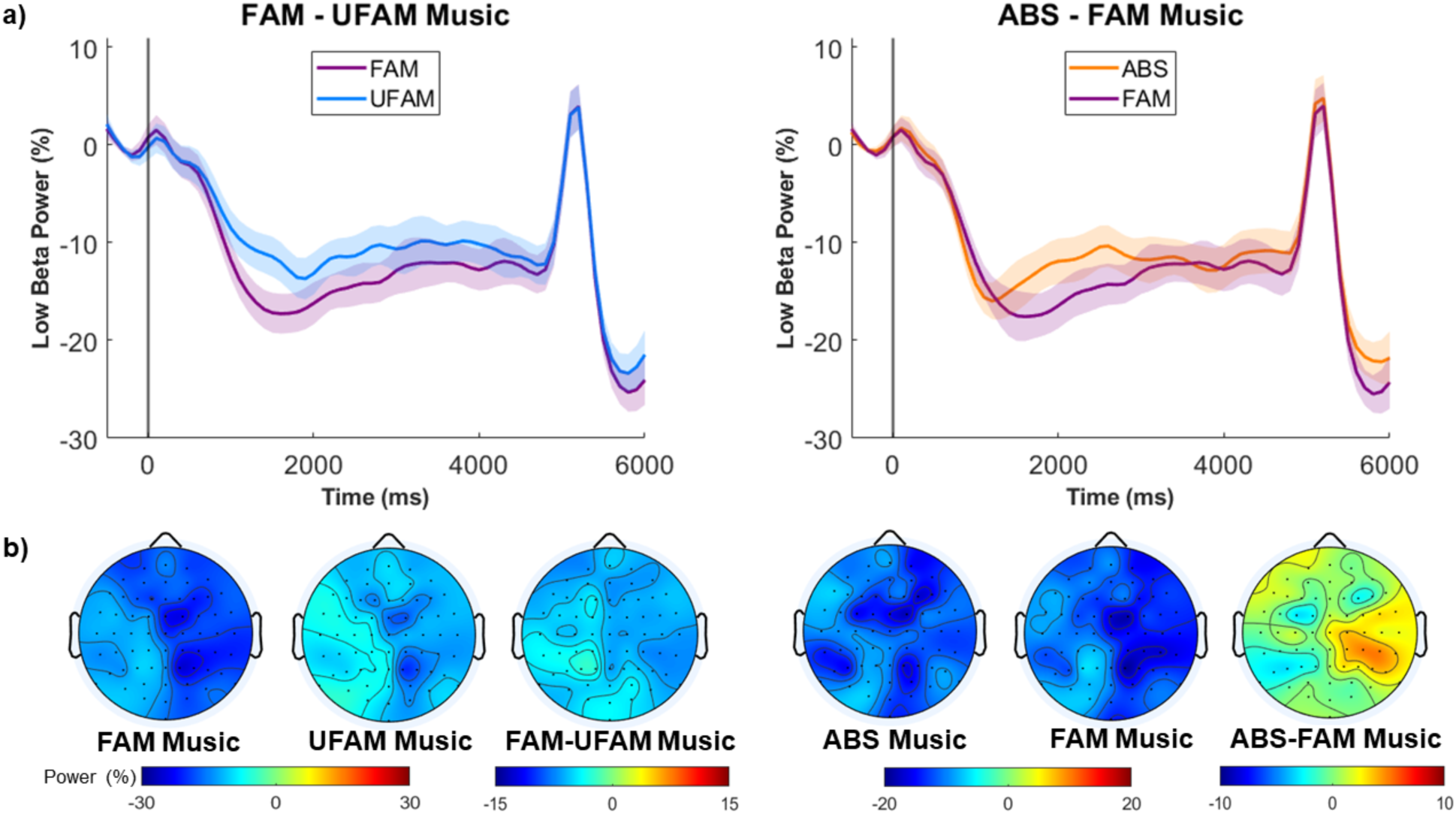
**a)** Changes in mean low-beta power (12 Hz to 16 Hz) relative to the pre-stimulus baseline, averaged over electrodes that are encompassed in each significant cluster (Table 4). Greater alpha suppression is observed for FAM music compared to (left panel) UFAM music and (right panel) ABS music. The amplitudes of each music condition are represented by different colors, where UFAM is blue, FAM is purple, and ABS is orange. Time 0 indicates the onset of the stimuli. Shading around each line represents the mean ± SEM. *ABS* = Autobiographically salient; *FAM* = Familiar; *UFAM* = Unfamiliar **b)** Topographic maps of the average of the electrodes in the corresponding cluster, around the peak latency (+/- 100 ms) for each pairwise contrast, including FAM-UFAM music (left panel) and ABS-FAM music (right panel).

**Table 4.**
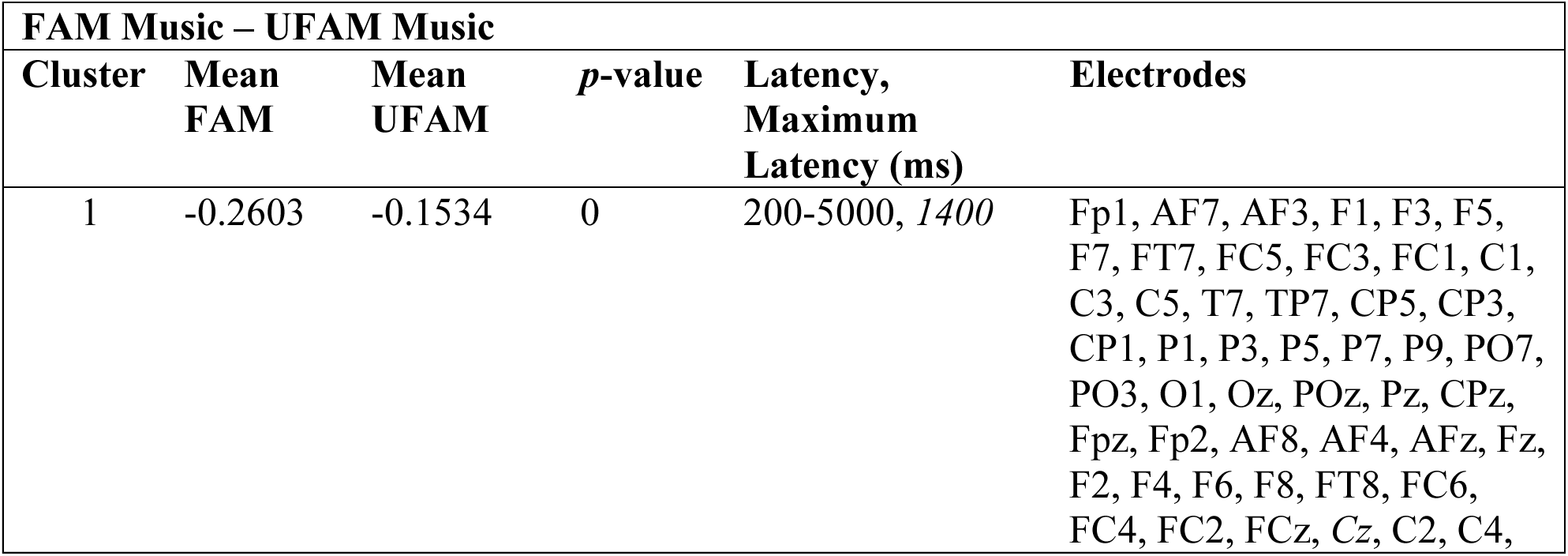
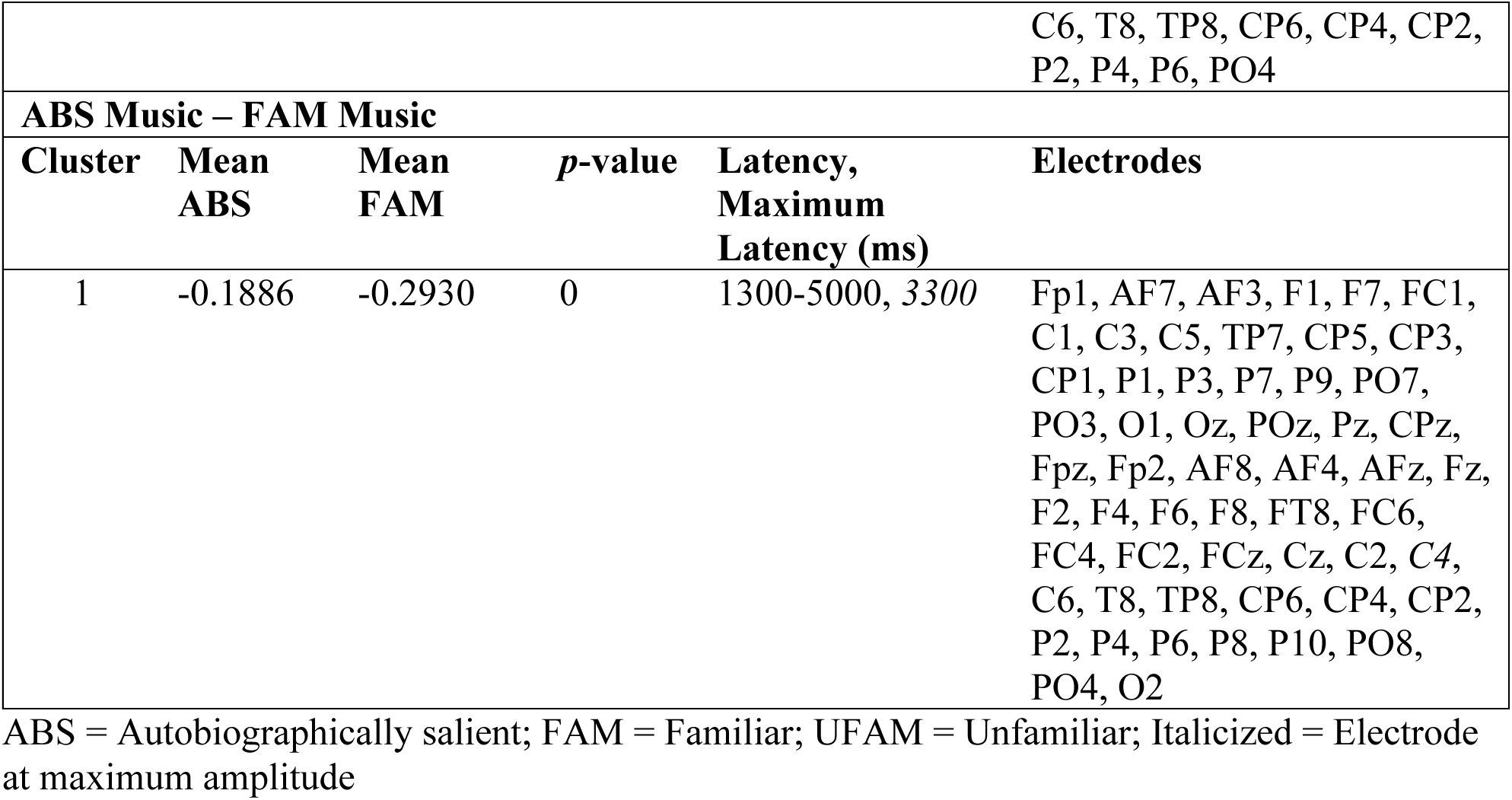
Summary of Cluster-Based Permutation Statistics from Low-Beta Band Power Analysis.

One cluster was revealed for the FAM-UFAM music contrast, whereby FAM showed greater low-beta power suppression compared to UFAM music. The cluster was widely distributed over bilateral frontal-central-parietal regions (Table 4) from 200 to 5000 ms, with the maximum difference occurring at 1400 ms (Fig 6a).

The ABS-FAM music contrast also revealed one significant cluster, indicating greater low-beta power suppression for FAM music relative to ABS music (*p* < 0.001). This bilateral cluster over frontal-central-parietal regions occurred later, between 1300 and 6000 ms, with the peak occurring at 2300 ms (Fig 6a).

#### Brain-Behavior Correlation Results

We explored brain-behavior relationships using ERP amplitude and band-specific frequency responses (i.e., alpha and low-beta) from 0 to 5000 ms and correlated these with RT, familiarity and memory ratings.

Cluster-based statistics revealed two significant associations between ERP amplitude and RT. First, a negative correlation (*p* = 0.001, *r* = -0.3637) was observed in bilateral parietal-occipital regions, indicating that greater negative amplitudes in this region were associated with faster RTs. In contrast, the second cluster was characterized by a positive relationship (*p* = 0.0026, *r* = 0.3538), linking larger amplitudes to slower RTs. However, this cluster was identified over left frontal regions with a later maximum latency. Stronger frontal activity may suggest greater cognitive control demands or response monitoring in trials where participants took more time to categorize music conditions from Experiment 1.

Alpha power was negatively correlated with memory ratings (*p* = 0.0464, *r* = -0.3491), indicating that lower alpha power is associated with higher memory ratings. Notably, this effect emerged earlier in time. A later cluster revealed a positive correlation between alpha power and RT (*p* = 0.0456, *r* = 0.3663), linking lower alpha power to faster identification of music excerpts. Taken together, the temporal sequence of these effects suggests that alpha suppression contributes to attentional engagement [43] and/or predictive processing [44] that facilitates faster perceptual identification.

A moderate negative association between low-beta power and familiarity ratings was revealed (*p* = 0.0426*, r* = -0.4024), indicating that lower beta power is related to higher familiarity ratings. Around the same time, a cluster characterized by a positive association was revealed between low-beta power and RT (*p* = 0.029, *r* = 0.4304). Similar to the pattern observed for alpha power, the temporal sequence suggests that reductions in low-beta power may reflect action preparation [45] due to familiarity-based processes that contribute to faster identification of music excerpts. For a detailed list of significant clusters identified in the brain-behavior correlation, see Table 5.

**Table 5.**
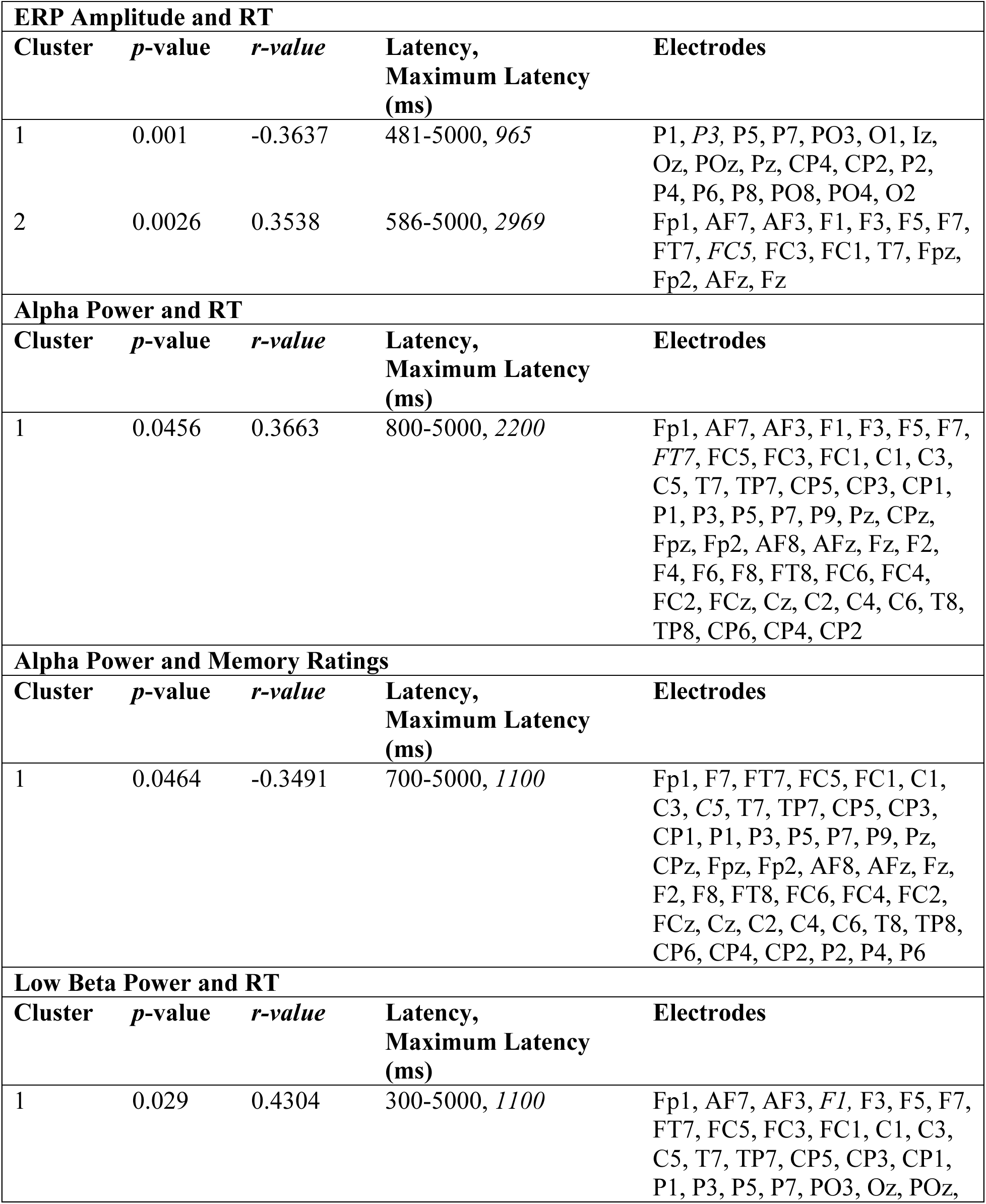
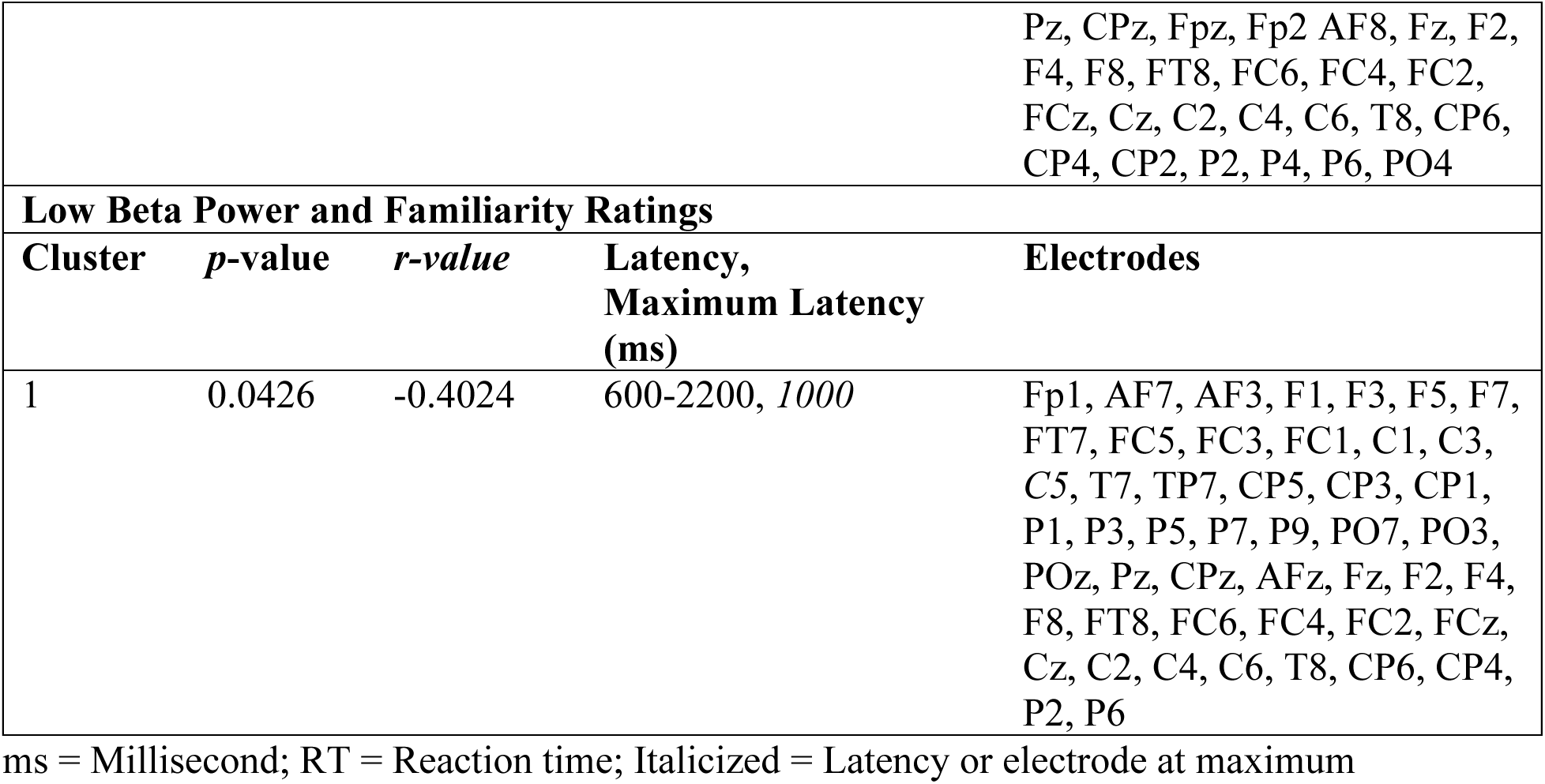
Summary of Cluster-Based Permutation Statistics from Brain-Behavior Correlation Analysis.

## Discussion

The present study investigated the behavioral and neural responses of older adults when listening to music excerpts that varied in personal significance. We were particularly interested in determining whether music associated with significant personal events (i.e., ABS music) generates brain responses that differ from FAM music. The first experiment showed that older adults were faster at identifying ABS music than FAM or UFAM music. On average, older adults took slightly over 2100 ms to correctly identify whether the incoming musical except was associated with an autobiographical event. We also observed a higher number of correct responses when identifying ABS than FAM and UFAM music. ERP analyses revealed larger ERP amplitudes over the right frontal-central areas in ABS than FAM and UFAM that began at 1457 ms and extended until the end of the epoch, which is in line with the LFE. Contrastingly, FAM music displayed an LPC relative to UFAM music. Time-frequency results further supported a distinction between ABS and FAM music. Compared to the latter, ABS music exhibited low-beta power suppression. When examining brain-behavior relationships, we found significant associations between ERP amplitude and RT and amplitude and memory ratings. In addition, alpha power was associated with RT and memory ratings, while low-beta power was correlated with RT and familiarity ratings.

### Electrophysiological Distinctions between Familiar and Autobiographically Salient Music

Our ERP findings show that listening to FAM and ABS music elicit distinct neural signatures. We hypothesized to observe an FN400 for the FAM-UFAM music contrast. Contrary to this, we found that FAM music generated higher amplitudes than UFAM over left parietal-occipital regions, similar to an LPC, which is posited to index recollection. Wilding and Rugg [27] found that the parietal ERP effect is related to the amount or quality of episodic detail during retrieval. This may indicate that participants in the present study had some level of episodic memory with the FAM music, albeit weaker than ABS music.

An alternative explanation for the LPC derives from more recent work that has challenged the traditional view, proposing instead that it reflects decision-dependent memory to guide judgment [46–48]. Ratcliffe et al. [47] used drift-diffusion model analysis to test whether the FN400 or LPC could predict drift rate, the rate at which evidence accumulates toward a decision. They found that only the LPC tracked memory strength for old words. Further support comes from Brezis and colleagues [46], who modified a Remember/Know paradigm by incorporating confidence judgments before asking participants to provide a “Remember”, “Know”, or “Guess” response. They found that LPC amplitude was larger for high-confidence “Know” responses than for low-confidence “Remember” responses, undermining the notion that the LPC is exclusive to recollection. In the context of our study, this LPC interpretation suggests that the observed effect reflect stronger memory signals driving more confident decision-making judgments.

While Malekmohammadi and colleagues [32] reported an old/new effect for FAM compared to UFAM over left frontal-central scalp regions from 400 to 450 ms, we observed the opposite effect, with FAM eliciting more negative ERP amplitude than UFAM music. This discrepancy may be related to the reference choice (i.e., average reference vs. linked mastoids) or other methodological factors including stimuli selection (e.g., self-chosen vs. experimenter-chosen, vocal vs. instrumental, and audio attribute comparability) and participant population (e.g., young adults vs. older adults).

It was hypothesized that an LPC would be observed for the ABS-UFAM music contrast, consistent with the late ERP cluster reported by Jagiello et al. [31]. In contrast, we found that listening to ABS compared to UFAM music generated a positive-going modulation over right frontal regions that began around 1450 ms and lasted until the end of the epoch. This cluster is topographically similar to an early cluster found by Jagiello and colleagues [31] but differs in latency and amplitude, which can be explained by study design, including sample size, baseline and epoch length, filtering, and whether the ABS and UFAM stimuli were measurably comparable in audio attributes. It is worth noting that the next cluster observed in our study, whereby ABS music was more positive than UFAM music, beginning around 580 ms over left parietal regions, is similar to the second cluster reported by Jagiello et al. [31].

We had also hypothesized that the ABS-FAM music contrast would elicit an LFE. Indeed, consistent with this hypothesis, we found that when older adults listened to ABS music compared to FAM music, an LFE emerged over right frontal-central regions. Aging studies have revealed that the LFE is more enhanced in older adults compared to young adults [49]. In the present study, the LFE effect was observed for a longer latency (600 to 2000 ms) than that typically cited in the literature, which may be related to our sample of older adults, who often exhibit slower cognitive processing speeds compared to young adults [50,51].

Three propositions have been put forth concerning the function of the LFE. One suggestion is that it indexes post-retrieval processing [27,28,52]. The second, is that the LFE indexes the retrieval of self-relevant information [29,30]. Lastly, the LFE has been associated with retrieval effort when memory representations are weak [53]. The latter seems unlikely given that participants provided specific song titles and names of artists which prompt a memory of a significant person, place, or event. Further, ABS music is deeply encoded, possesses a unique quality of involuntary retrieval and is emotionally rich. Therefore, our results are more in line with post-retrieval processing and/or self-relevant representations.

The present study revealed distinct patterns of spectral power modulation across conditions, particularly in the alpha (8 to 12 Hz) and low-beta (12 to 18 Hz) frequency ranges. Consistent with previous research [32], our contrasts between FAM and UFAM music indicated significant power differences in the alpha band, whereby greater suppression was observed for the former condition. Alpha power suppression is a well-established marker of enhanced auditory attention [43,54], including during music listening [32,44,55]. Thus, in the present study, the observed changes may reflect increased engagement and attention when listening to FAM music due to prior experience.

Another possible explanation for the observed event-related desynchronization of alpha power is that it may reflect predictive processes. Ito et al. [44] presented non-musicians with 20 FAM and 20 UFAM melodies, which were modified to include three seconds of silence in the middle of the excerpt. Participants were asked to predict the melody during the silent section. Time frequency analysis revealed that relative to UFAM melodies, FAM melodies exhibited greater alpha power suppression in left frontal-central regions, just before the silent section. This suggests that the brain engages in anticipatory processing when listening to FAM music, a mechanism that may also be at play in the present study.

In the current study, the maximum difference of the FAM-UFAM music contrast occurred over the left frontal-temporal scalp area (i.e., FT7 electrode), which broadly maps onto the left inferior frontal gyrus. Relatedly, this brain region was found to be one of the commonly activated areas when listening to FAM music as compared to UFAM music and may indicate prediction processes and/or subvocalization [17]. However, given the spatial limitations of EEG and the absence of additional imaging techniques, such as fMRI, this interpretation should be taken with caution.

While the FAM-UFAM music contrast primarily indicated differences in the alpha band, the ABS-FAM music was marked by changes in the low-beta band. Specifically, FAM music elicited greater suppression compared to ABS music peaking over the midline-right central scalp area (C2 electrode). Given that motor imagery can lead to event-related desynchronization over sensorimotor areas [45,56], this may suggest that FAM music evoked stronger, more automatic motor tendencies, such as an urge to tap or move along to the music, which participants actively suppressed during the task. The effect was observed over the right central scalp area, a site corresponding to sensorimotor areas including the supplementary motor cortex, which has been linked to FAM music listening [17]. A contributing factor to the observed effect is the frequency exposure of FAM compared to ABS music. While ABS music was selected by participants based on its significance and associations to important people, places, and events, it does not necessarily follow that these songs were frequently listened to in daily life. In contrast, FAM music consisted of widely popular songs that were often listed as a “hit” on the Billboards charts (see Table 1, Supplementary Material), and thus, frequently played on the radio or featured in films and other media. This increased exposure may have reinforced sensorimotor associations over time, leading to more tightly linked auditory-motor responses.

### Musical Memory: Single- or Dual-Process Model?

There is significant controversy regarding whether recognition memory is supported by one or more processes. Two dominant theoretical accounts have been proposed. The first is a single-process model, which describes recognition memory as a signal detection process, where memory strength exists along a continuum. Recognition judgments are based on the strength of an item’s memory signal, which reflects the degree to which the stimulus is similar to stored information [57]. Neural responses scale with memory strength, such that low recollection falls below a decision threshold, leading to weaker neural responses, and high recollection, exceeds the threshold, producing stronger neural responses. This model therefore predicts a graded effect, which would be present within the same scalp region [27,58]. Contrastingly, the dual-process model identifies temporally and topographically distinct modulations associated with familiarity and recollection [59]. According to this account, recognition memory involves an early sense of familiarity, as indexed by the FN400, followed by a later recollection response, marked by the LPC (for a review, see Yonelinas [60]). Lastly, although the late right frontal positivity effect has not been tied to any particular model, it is often associated with source memory decisions. Indeed, damage to the prefrontal cortex has been associated with impaired source memory performance [61,62].

Our findings reveal distinctions in neurophysiological activity when listening to music of varying levels of personal significance. Although our study bears some conceptual resemblance to the study-test paradigms often employed in recognition memory, the present design diverges in certain aspects. First, there were no additions of “new” stimuli. Second, confidence ratings are traditionally included in study-test paradigms to further distinguish between familiarity-based and recollection-based recognition, which were not collected in the present study. Future studies could incorporate confidence ratings to disentangle if the LPC observed in FAM music listening is related to decision making. Third, study-test paradigms do not typically include stimuli that are deeply relevant to the individual. Thus, the element of autobiographical memory and associated emotions may introduce a layer of complexity. Relatedly, the use of music, which evolves over time, may evoke neural responses that differ from those that arise from more static stimuli, such as symbols and images. Despite these differences in task design, we found strong evidence for distinct scalp distributions and latencies between FAM and ABS music, such as the LPC and the LFE, which point towards a model that involves more than one process in the context of musical memory.

### Limitations

We sought to investigate the manner in which ABS music engages memory processes by comparing this to FAM and UFAM music. Using a RT task, we first aimed to establish the time required for older adults to correctly identify music from each condition. Although we used 15 songs per condition and randomized the presentation of the clips to reduce the likelihood of a habitual response, one limitation was that each excerpt was repeated multiple times. Future studies could use excerpts that were clipped from various parts of the same song, while maintaining a large set of stimuli.

Music is a complex stimulus due to its many constituent features (i.e., rhythm, timbre, etc.), and the dynamic nature of one or more musical elements could influence the observed neural responses. Across conditions, our acoustic analysis of the stimuli showed no significant differences in RMS, tempo, or zero crossing rate. However, there were significant differences in onset rise time, pitch, spectral centroid, and spectral bandwidth. Nevertheless, low-level acoustic differences would be reflected in the early encoding portion of the evoked potential (i.e., N1, P2) and are, therefore, unlikely to have affected long latency responses.

Finally, we did not collect ratings of emotion during the task. As a result, we are unable to directly assess the extent to which emotional aspects of autobiographical remembering contributed to the observed neural responses. Notably, the ERPs for the ABS music were right-lateralized in frontal-central regions, a pattern that aligns with previous neuroimaging studies linking right-hemisphere activity to emotional aspects of autobiographical memory retrieval [63]. Further, given that our sample included older adults, in some cases, their ABS songs included music associated to loved ones who have passed away, suggesting that the stimuli may have evoked emotionally charged memories. However, without subjective measures of emotion, this interpretation remains speculative. More EEG research is needed to elucidate the timing and scalp distribution of emotional processes when listening to ABS songs, including distinctions among emotions such as happiness, sadness, and more complex emotions, such as nostalgia.

## Conclusion

Our behavioral and electrophysiological data provide converging evidence that ABS music uniquely engages memory-related processes. Specifically, ABS music elicits quicker RTs and distinct neural signatures than FAM music. The ERP elicited by ABS music revealed a sustained late right frontal positivity, whereas listening to FAM music was associated with an early LPC over the left posterior-occipital regions. Time-frequency analyses revealed less low-beta suppression for ABS music compared to FAM music, while conversely, FAM music showed greater alpha suppression compared to UFAM music. Brain-behavior correlations further highlight relationships between ERP amplitude and both RT and memory ratings. Furthermore, alpha power was correlated with both RT and memory ratings, while beta power showed associations with RT and familiarity ratings. Taken together, the temporal, spectral, and topographic differences between FAM and ABS music add to the evidence that distinct cognitive and neural processes underpin familiarity and deeper recollection. These findings underscore the need to distinguish between the two conditions in musical memory studies and offer meaningful implications for ABS music-based interventions in aging and memory-related disorders, such as dementia.

## Acknowledgements

Thank you to Ashna Imran, Madison W. Grassi, and Shimin Mo for assistance in preparing the auditory stimuli and/or data collection. Finally, we thank all those who participated in the experiments reported here.

## Author Contributions

Veronica Vuong (Conceptualization, Data curation, Formal analysis, Investigation, Methodology, Project administration, Software, Visualization, Writing – original draft), Mary O’Neil (Data curation, Investigation), Andrew Dimitrijevic (Methodology, Writing – review & editing), Bradley R. Buchsbaum (Writing – reviewing & editing), Michael H. Thaut (Conceptualization, Methodology, Funding acquisition, Supervision, Validation, Writing – reviewing & editing), Claude Alain (Conceptualization, Methodology, Funding acquisition, Resources, Supervision, Validation, Writing – review & editing).

