## Supplementary Table 1 for "When Melodies Cue Memories: Electrophysiological Correlates of Autobiographically Salient Music Listening in Older Adults"

[1] Rotman Research Institute, Baycrest Academy for Research and Education, North York, ON, Canada
[2] Institute of Medical Science, Temerty Faculty of Medicine, University of Toronto, Toronto, ON, Canada
[3] Music and Health Science Research Collaboratory, Faculty of Music, University of Toronto, Toronto, ON, Canada
[4] Department of Psychology, Faculty of Health, York University, Toronto, ON, Canada
[5] Department of Otolaryngology, Head and Neck Surgery, Sunnybrook Health Sciences Centre, Toronto, ON, Canada
[6] Department of Otolaryngology, Head and Neck Surgery, Temerty Faculty of Medicine, University of Toronto, Toronto, ON, Canada
[7] Evaluative Clinical Sciences Platform, Sunnybrook Research Institute, Toronto, ON, Canada
[8] Department of Physiology, University of Toronto, Toronto, ON, Canada
[9] Department of Psychology, University of Toronto, Toronto, ON, Canada
[10] Rehabilitation Sciences Institute, Temerty Faculty of Medicine, University of Toronto, Toronto, ON, Canada
[‡] Co-Senior Authors

**Corresponding author:**
**Veronica Vuong**

**Table 1: Examples of Participant Stimuli**

| # | ABS Song (Title, Artist, [Year]) | FAM Song (Title, Artist, [Year]) | # Streams* | Billboard Chart | UFAM Song (Title, Artist, [Year]) | # Streams* | Billboard Chart |
|---|---|---|---|---|---|---|---|
| 1 | Windy – The Association (1967) | Happy Together – The Turtles (1967) | 449 million | Hot 100 (1967) | Come Round Here – The Cowsills (1967) | 736,400 | - |
| 2 | Hey! Bo Diddley – Bo Diddley (1957) | Johnny B. Goode – Chuck Berry (1958) | 376 million | Hot 100 (1958) | All Night Boogie (All Night Longer) – Howlin' Wolf (1960) | 593,000 | - |
| 3 | A Whiter Shade of Pale – Procol Harum (1967) | Nights in White Satin – The Moody Blues (1967) | 238 million | Hot 100 (1964) | Day of the Fool – Aphrodite's Child (1969) | 40,600 | - |
| 4 | Stand By Me – Ben E. King (1962) | Under the Boardwalk – The Drifters (1964) | 209 million | Hot 100 (1964) | So Many Cute Little Girls – Jackie Wilson (1960) | 9,900 | - |
| 5 | Made You Look – Meghan Trainor (2022) | About Damn Time – Lizzo (2022) | 729 million | Hot 100 (2022) | I Can Be Your Man – Betty Who (2022) | 370,000 | - |
| 6 | Big Girls Don't Cry – Frankie Valli & The Four Seasons (1962) | My Girl – The Temptations (1964) | 463 million | Hot 100 (1965) | In that Great Gettin' Up Mornin' – The Righteous Brothers (1962) | 16,848 | - |
| 7 | Respect – Aretha Franklin (1967) | I Heard It Through the Grapevine – Marvin Gaye (1968) | 383 million | Hot 100 (1968) | Just Look What You've Done – Brenda Holloway (1967) | 619,870 | Hot 100 (1967) |
| 8 | Up on the Roof – The Nylons (1981) | You Make My Dreams Come True – Hall & Oates (1980) | 881 million | Hot 100 (1981) | When You're Young and in Love – The Flying Pickets (1983) | 1.1 million | - |
| 9 | A Hard Day's Night – The Beatles (1964) | I'm A Believer – The Monkees (1966) | 386 million | Hot 100 (1967) | Painter Man – The Creation (1966) | 678,000 | - |
| 10 | Kodachrome – Paul Simon (1973) | Country Roads – John Denver (1971) | 732 million | Hot 100 (1971) | Man in the Mirror – Graham Nash (1971) | 650,000 | - |
| 11 | Diamonds on the Soles of Her Shoes – Paul Simon (1986) | Hungry Eyes – Eric Carmen (1987) | 606 million | Hot 100 (1988) | It's Alright – Chicago (1986) | 189,000 | - |
| 12 | All I Have to do is Dream – Everly Brothers (1958) | We Belong Together – Ritchie Valens (1958) | 110 million | - | The Loneliest Sound – Ricky Nelson (1959) | 9,400 | - |
| 13 | The Lion Sleeps Tonight – The Tokens (1961) | Runaround Sue – Dion (1961) | 185 million | Hot 100 (1961) | The Diary – Little Anthony & The Imperials (1959) | 357,015 | - |
| 14 | Fire and Rain – James Taylor (1970) | Landslide – Fleetwood Mac (1975) | 662 million | Hot 100 (1998) | Nine Houses – Seals & Crofts (1973) | 118,000 | - |
| 15 | Calm – Rema (2022) | Buga – Kizz Daniel (2022) | 229 million | Year-End U.S. Afrobeats (2022) | Cruise – SPINALL (2022) | 299,000 | - |
| 1 | Don't Worry, Be Happy – Bobby McFerrin (1988) | Buffalo Solider – Bob Marley and the Wailers (1983) | 636 million | Hot 100 (1983) | Sheila – Gregory Issacs (1983) | 210,000 | - |
| 2 | I Will Always Love You – Whitney Houston (1992) | Save the Best for Last – Vanessa Williams (1991) | 201 million | Hot 100 (1992) | Better Off – Vanessa Williams (1991) | 96,000 | - |

| | | | | | | | |
|---|---|---|---|---|---|---|---|
| 3 | I Will Never Find Another You – The Seekers (1964) | California Dreamin' – The Mamas and The Papas (1965) | 987 million | Hot 100 (1966) | That's the Way I Feel – Crispian St. Peters (1966) | 48,500 | - |
| 4 | Imagine – John Lennon (1971) | Time in a Bottle – Jim Croce (1972) | 233 million | Hot 100 (1973) | Be Nice to Me – Todd Rundgren (1971) | 739,000 | - |
| 5 | Hello – Lionel Richie (1983) | True – Spandau Ballet (1983) | 473 million | Hot 100 (1983) | Love's Been Here and Gone – James Ingram (1986) | 71,129 | - |
| 6 | My Heart Will Go On – Céline Dion (1997) | Angel – Sarah McLachlan (1995) | 232 million | Hot 100 (1999) | Leading With Your Heart – Barbra Streisand (1997) | 141,553 | - |
| 7 | Yesterday – The Beatles (1965) | Happy Together – The Turtles (1967) | 449 million | Hot 100 (1967) | I've Got That Feeling – The Kinks (1964) | 396,000 | - |
| 8 | The End of the World – Skeeter Davis (1962) | Crazy – Patsy Cline (1961) | 141 million | Hot 100 (1962) | Take Me as I Am – Don Gibson (1958) | 77,429 | - |
| 9 | River of No Return – Tennessee Ernie Ford (1957) | That's Amore – Dean Martin (1953) | 193 million | Hot 100 (1953) | Losing Your Love – Jim Reeves (1961) | 759,000 | - |
| 10 | Singin' In the Rain – Gene Kelly (1952) | Banana Boat (Day-O) – Harry Belafonte (1956) | 123 million | Hot 100 (1957) | Sisters – Peggy Lee (1954) | 267,448 | - |
| 11 | Candle In the Wind – Elton John (1973) | American Pie – Don McLean (1971) | 643 million | Hot 100 (1972) | Gently I'll Wake You – Chicago (1976) | 393,000 | - |
| 12 | Unchained Melody – The Righteous Brothers (1965) | My Girl – The Temptations (1965) | 798 million | Hot 100 (1965) | If Mary's There – Gene Pitney (1965) | 18,549 | |
| 13 | Man in the Mirror – Michael Jackson (1988) | Purple Rain – Prince (1984) | 586 million | Hot 100 (1984) | I Really Love You – Bobby Brown (1988) | 807,845 | - |
| 14 | Forever and For Always – Shania Twain (2002) | Breathe – Faith Hill (1999) | 151 million | Hot 100 (1999) | Just Another Heartache – Chely Wright (1997) | 238,400 | - |
| 15 | Always On My Mind – Elvis Presley (1972) | If You Could Read My Mind – Gordon Lightfoot (1972) | 126 million | Hot 100 (1971) | (Last Night) I Heard You Crying in Your Sleep – Roy Orbison (1970) | 95,099 | - |
| 1 | Year of the Cat – Al Stewart (1976) | Hotel California – The Eagles (1976) | 1.4 billion | Hot 100 (1977) | Gently I'll Wake You – Chicago (1976) | 391,000 | - |
| 2 | Full Circle – Half Moon Run (2013) | Counting Stars – OneRepublic (2013) | 5.7 billion | Hot 100 (2014) | Byebyelove – Jimmy Eat World (2013) | 952,000 | - |
| 3 | Take On Me – A-Ha (1985) | Don't You (Forget About Me) – Simple Minds (1985) | 1 billion | Hot 100 (1985) | Waiting For You – Corey Hart (1985) | 78,000 | - |
| 4 | Hungry Heart – Bruce Springsteen (1980) | Jack & Diane – John Mellencamp (1982) | 352 million | Hot 100 (1982) | Red Hot & Blue Love – Rick Springfield (1980) | 161,300 | - |
| 5 | Heroes – David Bowie (1977) | We Are the Champions – Queen (1977) | 862 million | Hot 100 (1978) | Tried to Love – Peter Frampton (1977) | 111,000 | Hot 100 (1978) |
| 6 | She Loves You – The Beatles (1964) | Louie, Louie – The Kingsmen (1963) | 138 million | Hot 100 (1963) | What Kind of Boy – The Hollies (1964) | 107,159 | - |
| 7 | Love Over and Over – The McGarrigles (1982) | Everywhere – Fleetwood Mac (1987) | 700 million | Hot 100 (1988) | I'm Dancing as Fast as I Can – Juice Newton (1982) | 87,000 | - |

| | | | | | | | |
|---|---|---|---|---|---|---|---|
| 8 | These Foolish Things – Bryan Ferry (1973) | Me and Julio Down by The Schoolyard – Simon & Garfunkel (1972) | 291 million | Hot 100 (1972) | Nine Houses – Seals & Crofts (1973) | 118,000 | - |
| 9 | Another Day – Paul McCartney (1971) | Mrs. Robinson – Simon & Garfunkel (1968) | 583 million | Hot 100 (1968) | I'll Be the One – Badfinger (1971) | 447,000 | - |
| 10 | You Baby – The Mamas and The Papas (1966) | Good Vibrations – The Beach Boys (1966) | 365 million | Hot 100 (1966) | The Lover – Tommy James (1966) | 63,900 | - |
| 11 | Take Me to Church – Hozier (2013) | Set Fire to the Rain – Adele (2011) | 1.2 billion | Hot 100 (2012) | Okay – Winter Aid (2017) | 517,309 | - |
| 12 | When God Made Me – Neil Young (2005) | Drops of Jupiter – Train (2001) | 329 million | Hot 100 (1983) | God's Golden Eyes – John Hiat (2000) | 114,442 | - |
| 13 | Over You – Roxy Music (1980) | Love Will Tear Us Apart – Joy Division (1980) | 423 million | Disco Top 100 (1980) | I Think I'm Go Go – Squeeze (1980) | 187,700 | - |
| 14 | Everywhere – Lucy Wainwright Roche (2007) | Waiting on the World to Change – John Mayer (2006) | 409 million | Hot 100 (2006) | Anna – Antje Duvekot (2006) | 69,230 | - |
| 15 | A New England – Billy Bragg (1983) | Every Breath You Take – The Police (1983) | 2.6 billion | Hot 100 (1983) | The Time Is Now – Naked Eyes (1983) | 62,700 | - |
| 1 | My Way – Frank Sinatra (1969) | Sway – Dean Martin (1964) | 262 million | - | A Little Voice – Dean Martin (1964) | 234,000 | - |
| 2 | Crocodile Rock – Elton John (1973) | You Make Me Feel Like Dancin' – Leo Sayer (1976) | 664 million | Hot 100 (1977) | Don't Start Me to Talkin' – The Doobie Brothers (1973) | 645,000 | - |
| 3 | You're So Vain – Carly Simon (1972) | It's Too Late – Carole King (1971) | 197 million | Hot 100 (1971) | No. 1 (Ahwantgemmeh) – Melissa Manchester (1974) | 3,458 | - |
| 4 | I Will Survive – Gloria Gaynor (1978) | Got to be Real – Cheryl Lynn (1978) | 105 million | Hot 100 (1979) | You Touch My Hot Line – The Trammps (1976) | 167,272 | - |
| 5 | Hotel California – The Eagles (1976) | Another Brick in The Wall, Part 2 – Pink Floyd (1979) | 753 million | Hot 100 (1980) | Turn It Loose – The Doobie Brothers (1976) | 561,316 | - |
| 6 | Sweet Home Alabama – Lynyrd Skynyrd (1974) | The Joker – Steve Miller Band (1973) | 418 million | Hot 100 (1974) | Alta Mira – The Edgar Winter Group (1972) | 369,859 | - |
| 7 | Colour My World – Chicago (1970) | Let It Be – The Beatles (1970) | 700 million | Hot 100 (1970) | Only a Moment Ago – The Partridge Family (1970) | 542,308 | - |
| 8 | Ordinary Miracles – Amy Sky (1998) | Because You Loved Me – Céline Dion (1996) | 605 million | Hot 100 (1996) | In My Arms Again – Michael W. Smith (1998) | 215,069 | - |
| 9 | Up! – Shania Twain (2003) | This Kiss – Faith Hill (1998) | 156 million | Hot 100 (1998) | Feel It Comin' On – Sara Evans (2003) | 185,455 | - |
| 10 | Brown Eyed Girl – Van Morrison (1967) | I'm a Believer – The Monkees (1967) | 386 million | Hot 100 (1966) | Too Young to Be the One – The Turtles (1967) | 686,292 | - |
| 11 | Jolene – Dolly Parton (1973) | If You Could Read My Mind – Gordon Lightfoot (1970) | 118 million | Hot 100 (1971) | What a Man My Man Is – Lynn Anderson (1974) | 176,056 | - |
| 12 | Could You Be Loved – Bob Marley and the Wailers (1980) | Red, Red Wine – UB40 (1983) | 705 million | Hot 100 (1988) | Any Day Now – Dennis Brown (1982) | 327,839 | - |

| | | | | | | | |
|---|---|---|---|---|---|---|---|
| 13 | Leaving on a Jet Plane – John Denver (1966) | Our House – Crosby, Stills, Nash & Young (1970) | 137 million | Hot 100 (1970) | Flying – Badfinger (1970) | 459,474 | - |
| 14 | Downtown – Petulia Clark (1964) | Rhythm of the Rain – The Cascades (1963) | 71 million | Hot 100 (1963); Easy Listening (1963) | More – Vic Dana (1966) | 2.2 million | - |
| 15 | Stairway to Heaven – Led Zeppelin (1971) | All Along the Watchtower – Jimi Hendrix (1968) | 853 million | Hot 100 (1968) | Inside the Keeper's Pantry – Bill Fay (1971) | 61,003 | - |

**Encompasses the combined number of streams on Spotify and Youtube recorded at the time of study. ABS = Autobiographically salient; FAM = Familiar; UFAM = Unfamiliar*
